# Ambient RNA analysis reveals misinterpreted and masked cell types in brain single-nuclei datasets

**DOI:** 10.1101/2022.03.09.483658

**Authors:** Emre Caglayan, Yuxiang Liu, Genevieve Konopka

**Author notes:** Corresponding authors: Emre Caglayan, Genevieve Konopka.

## Abstract

Ambient RNA contamination in single-cell RNA sequencing (RNA-seq) is a significant problem, but its consequences are poorly understood. Here, we show that ambient RNAs in brain single-nuclei RNA-seq can have a nuclear or extra-nuclear origin with distinct gene set signatures. Both ambient RNA signatures are predominantly neuronal and we find that some previously annotated neuronal cell types are distinguished by ambient RNA. Strikingly, we also detect pervasive neuronal ambient RNA contamination in all glial cell types unless glia and neurons are physically separated prior to sequencing. We demonstrate that this contamination can be removed *in silico*. We also show that previous annotations of immature oligodendrocytes are likely glial cells contaminated with ambient RNAs. After ambient RNA removal, we detect extremely rare, committed oligodendrocyte progenitor cells, which were infrequently annotated in previous adult human brain datasets. Together, these results provide an in-depth analysis of ambient RNA contamination in brain single-cell datasets.

## Introduction

Single-cell RNA sequencing utilizes abundantly more cell barcodes than the number of targeted cells to ensure single-cell capture per droplet^1^. This results in a relatively small number of droplets that capture real cells as most droplets are ‘empty droplets’ and do not contain real cells^1^. However, ambient material created during sample processing can also contain transcripts called ‘ambient RNA’ which can be captured by empty droplets, making them appear non-empty in data analysis^2^. As a result, separation of empty droplets from real cells becomes a significant problem in single-cell RNA-seq analysis. Since real cells/nuclei contain more transcripts (and more unique molecular identifiers -UMIs- that denote unique reads) than ambient RNAs captured in empty droplets, it is possible to apply a cutoff based on number of UMIs to retain real cells. However, this can be inaccurate as UMI distribution of cell barcodes is not completely discrete which makes a hard cutoff arbitrary. Moreover, certain cell types may be transcriptomically more silent than others which could lead to filtering out by a UMI-based cutoff^3^. Recent tools addressed this problem by distinguishing real cells and empty droplets by using other metrics such as expression profile and nuclear fraction^2,4,5^.

In addition to the cell barcodes not associated with real cells, ambient RNAs can also be captured in the same droplet along with real cells resulting in ambient RNA contamination. This requires removal of the ambient RNA signature from these cell barcodes before further analysis. However, many previous studies have likely not removed ambient RNA contamination, therefore it is unknown whether ambient RNA contamination constitutes an important problem that leads to misinterpretations in the downstream analysis. Since an ambient RNA profile is primarily from the most abundant cell types, less abundant cell types may be especially more susceptible to misinterpretations. There is an array of recent tools that are specifically designed to remove ambient RNA contamination from real cells which can be utilized to assess the effect of ambient RNA contamination in single-cell analysis^4,6–9^. While these tools aim to be effective across different tissues, genetic signatures of ambient RNA contamination differ between tissue types. Thus, it is important to know which genes are overrepresented in ambient RNAs. This will aid both interpretation of previous datasets as well as assessment of ambient RNA contamination removal tools. In this study, we focus on single-nuclei RNA-seq (snRNA-seq) studies from brain tissue^10–13^. Brain datasets are also a good model to understand the effects of ambient RNA contamination since neurons are more abundant than any other cell type alone and contain more transcripts than glia in single-cell studies from adult mammalian cortex^14^. Therefore, ambient RNA profiles from single-cell studies of brain should be biased for neurons. We thus hypothesize that this bias may contribute to both empty droplets with neuronal signatures as well as distinctive neuronal read contamination in other cell types, mainly in glia.

Here, we analyzed ambient RNA signatures from human brain snRNA-seq datasets **(Supplementary Table 1)** by retaining additional cell barcodes that are typically removed due to low UMIs. We found two types of ambient RNAs separated by their intronic read ratio: extra-nuclear ambient RNAs with low intronic reads and nuclear ambient RNAs with high intronic reads. Comparisons with nuclei-sorted datasets revealed that extranuclear ambient RNA can be cleared by physical nuclei sorting. We show that the ambient RNA signature is predominantly neuronal and all glia are contaminated with ambient RNAs. Ambient RNA contamination in glia was removed in datasets depleted of neurons (NeuN-sorted) before droplet capture^15^. As an *in-silico* alternative, we used CellBender^9^ which also removed detectable ambient RNA contamination from glia. Finally, re-analysis of the oligodendrocyte lineage trajectory after ambient RNA removal revealed that previous annotations of immature oligodendrocytes are likely ambient RNA contaminated glia. Instead, we found a rare, transient cell type named COPs (committed oligodendrocyte progenitor cells). COPs were previously described in an adolescent mouse dataset^16^ but not described in most human snRNA-seq datasets with few exceptions^13,17^. Together, our results provide an in-depth analysis of ambient RNA contamination in single-nuclei brain datasets and reveal misinterpreted results that can be explained by ambient RNA contamination.

## Results

### Both nuclear and extra-nuclear ambient RNAs confound cell type annotation

Single-cell studies of adult brain tissue have repeatedly shown greater number of transcripts present in neurons compared to glia^14^. Interestingly, several snRNA-seq studies reported some neuronal cell types (e.g. Neu-NRGN and Neu-mat in **Figure 1A**) that have fewer transcript counts than other neurons^12,14^. We also noticed that Neu-NRGNs had higher mitochondrial reads than other neuronal cell types in a non-sorted dataset (referred to as NSD1^12^) **(Figure 1B)**. Considering the possibility of extra-nuclear transcript contamination in nuclei-based sequencing, cell barcodes with high extra-nuclear contamination should contain lower intronic read ratios since intronic reads will not be present in extra-nuclear transcripts. We thus hypothesized that Neu-NRGNs and Neu-mat may be represented by cell barcodes with high amount of such extra-nuclear transcript contamination. To test this hypothesis, we calculated the intronic read ratio per cell barcode (see methods) and found that Neu-NRGN but not Neu-mat displayed markedly lower intronic read ratios compared to other cell types **(Figure 1C)**. To test whether these cell types are similar to normally discarded cell barcodes with low UMI, we then clustered an excess number of cell barcodes that contained low UMIs together with the annotated cell barcodes in the original publication. To identify annotated cell barcodes that cluster with low UMI cell barcodes, we focused on the clusters that were largely, but not fully filtered out (>75% filtered) in the original publication and named them “ambient clusters” **(Figure 1D–E)**. We found that Neu-NRGN cell barcodes predominantly clustered with the ambient clusters while other cell types were almost absent in ambient clusters, except for Neu-mat **(Figure 1F)**. If Neu-NRGN cell barcodes are indeed highly contaminated with extra-nuclear ambient RNA, they should also be depleted of long non-coding RNAs (lncRNAs), which are retained in the nucleus^18^. Indeed, Neu-NRGN barcodes displayed fewer lncRNAs than other neurons **(Figure 1G)**. An interesting example was *MALAT1,* which has elevated expression in the brain^19^ and was depleted in Neu-NRGN cell barcodes **(Figure 1H)**. Together these results indicate that Neu-NRGN cell barcodes contain high extra-nuclear ambient RNA contamination.

**Figure 1.**
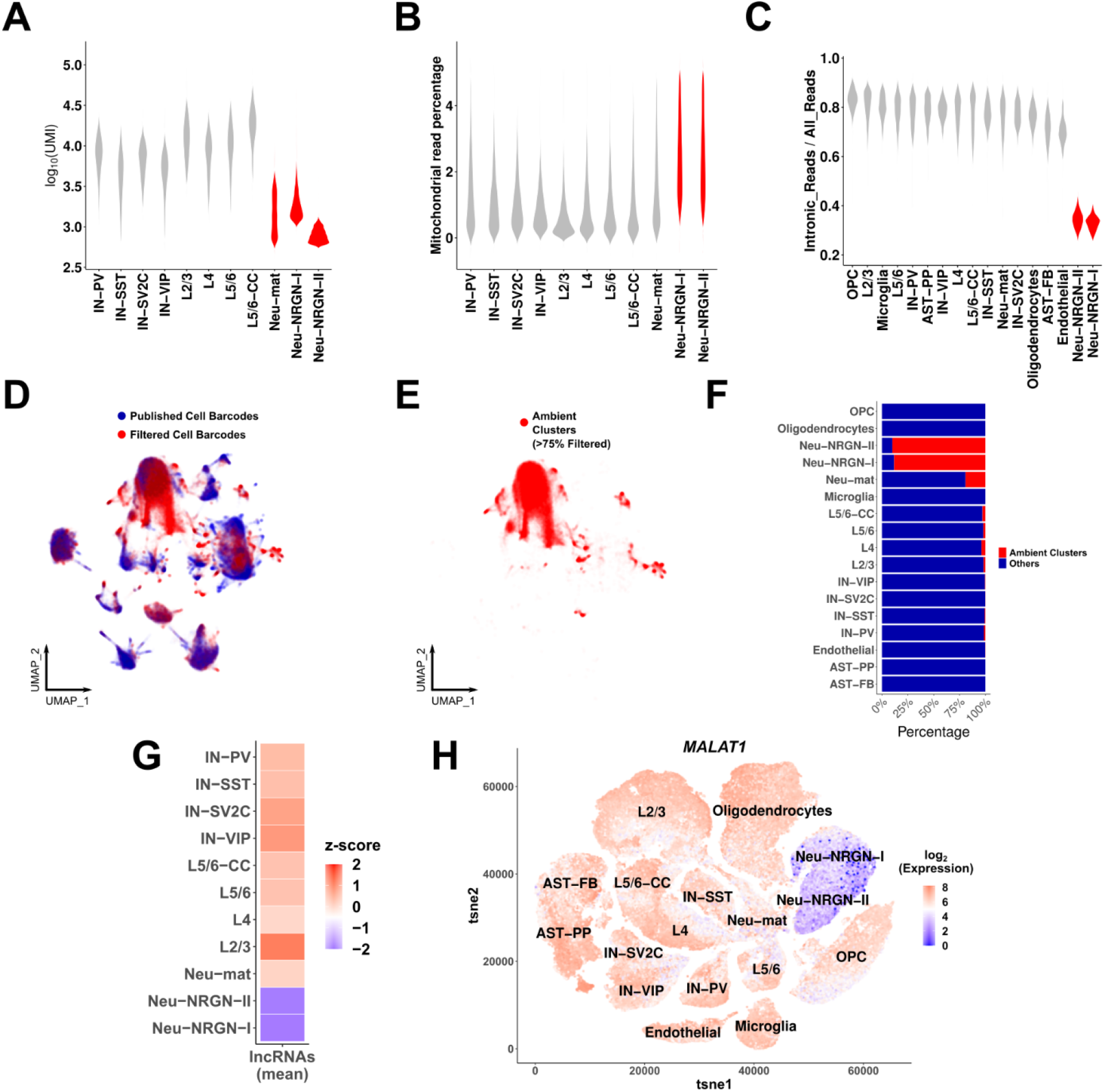
Neu-NRGNs are comprised of extra-nuclear ambient RNAs. **(A)** Log_10_ transformed UMI counts per neuronal cell type in NSD1 (non-sorted dataset 1). Cell types with unusually low UMI count are shown in red. **(B)** Mitochondrial read percentage per neuronal cell type in NSD1. Cell types with significantly high mitochondrial read percentage are shown in red (Wilcoxon rank sum test. P-value < 0.05). **(C)** Intronic read ratio of all cell types in NSD1. **(D)** UMAP representation after co-embedding of same numbers of published (blue) and filtered cell barcodes (red) (dataset: NSD1). **(E)** Clusters that are >75% composed of filtered cell barcodes are highlighted and named ambient clusters (dataset: NSD1). **(F)** Percentage of cell barcodes in ambient clusters per cell type. Red: ambient clusters, blue: other clusters. **(G)** Normalized, log-transformed and z-scored expression levels of lncRNAs across cell types. Mean of all lncRNAs were taken before z-score. **(H)** Normalized and log transformed expression level of *MALAT1* in all cells on the tSNE space.

We then hypothesized that extra-nuclear ambient RNA can be removed by fluorescence activated cell sorting (FACS) after nuclei isolation. To test this, we analyzed another cortical snRNA-seq dataset, named sorted dataset 1 (SD1) which performed nuclei sorting (purification of DAPI+ nuclei with flow cytometry)^20^. In contrast, the previous dataset (NSD1) performed nuclei isolation but not nuclei sorting by flow cytometry (Figure 1). Since low intronic read ratio indicates the presence of extra-nuclear transcripts, we then checked intronic read ratio in both datasets to assess whether extra-nuclear transcripts are removed by nuclei sorting. As expected, NSD1 displayed low intronic read ratio in cell barcodes with low UMI since these cell barcodes mostly carry ambient RNAs **(Figure 2A)**. In contrast, we found that intronic read ratio did not markedly change with increasing UMI in SD1 **(Figure 2B)**. We then assessed ambient RNA signatures after the removal of extra-nuclear ambient RNA and found ambient clusters in SD1 **(Figure 2C–D)**. Interestingly, SD1 ambient cluster markers were highly and distinctly enriched in the Neu-mat markers from NSD1 **(Figure 2E)**. We also observed significant enrichment of SD1 ambient cluster markers in other neuronal cell type markers although it was markedly less compared to Neu-mat. Co-clustering of SD1 ambient clusters and NSD1 cell types also grouped SD1 ambient clusters and Neu-mat cell barcodes together **(Figure 2F)**. Overall, these results indicate that Neu-mat cell barcodes carry an unusually high nuclear ambient RNA signature and Neu-NRGN cell barcodes are distinctly contaminated with extra-nuclear ambient RNA. These results also reveal that nuclei sorting, but not nuclei isolation, effectively removes extra-nuclear ambient RNA; however, it cannot remove nuclear ambient RNA. Therefore, we named NSD1 ambient markers as ‘extra-nuclear ambient markers’, and SD1 ambient markers as ‘nuclear ambient markers’ in the remainder of this study **(Supplementary Table 2)**.

**Figure 2.**
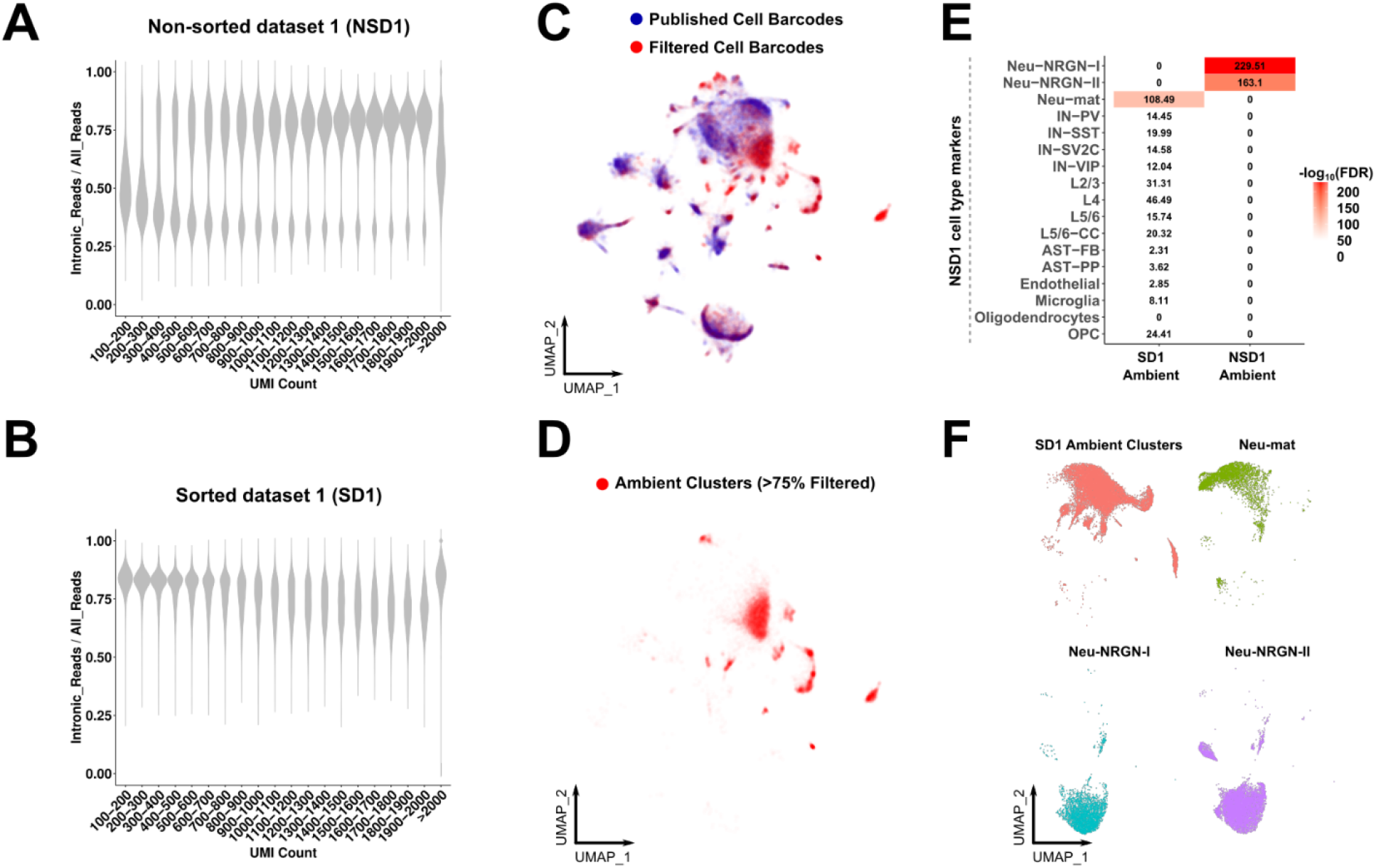
Extra-nuclear and nuclear ambient RNAs are distinct from each other. **(A-B)** Intronic read ratio across increasing UMI count in **(A)** non-sorted dataset 1 (NSD1) and **(B)** sorted dataset 1 (SD1). UMIs are divided into intervals of 100 from 100-2000. **(C)** UMAP representation after co-embedding of the same numbers of published (blue) and filtered cell barcodes (red) (dataset:SD1). **(D)** Clusters that are >75% composed of filtered cell barcodes are highlighted and named ambient clusters (dataset: SD1). **(E)** Enrichments between ambient RNA markers and Neu-mat or Neu-NRGN cell types in NSD1 (Fisher’s exact test, numbers correspond to log_10_(FDR)). **(F)** Co-embedding of Neu-NRGNs, Neu-mat and SD1 ambient clusters.

To provide independent support for these results, we then analyzed a second snRNA-seq cortical human dataset that did not include nuclei sorting (non-sorted dataset 2, NSD2) and as well as another cortical human dataset that did perform nuclei sorting (sorted dataset 2, SD2)^21^. We reproduced a similar pattern of increasing intronic read ratio with increasing UMI in the NSD2 whereas there was no marked difference in the SD2 **(Extended Figure 1A–B)**. We then similarly identified ambient clusters from the NSD2 dataset **(Extended Figure 1C)**. Intronic read ratio distribution in ambient clusters was bimodal and we divided them into two categories: cell barcodes with high intronic read ratio (High-Intron-CB) and cell barcodes with low intronic read ratio (Low-Intron-CB) **(Extended Figure 1D)**. In line with the previous results, genes overrepresented in Low-Intron-CB were highly enriched in extra-nuclear ambient RNA markers whereas High-Intron-CB were enriched in nuclear ambient RNA markers **(Extended Figure 1E)**.

### Signatures and sources of extra-nuclear and nuclear ambient RNAs

We then performed gene ontology enrichment on both ambient RNA types and found that extra-nuclear ambient RNA markers were enriched for genes involved in ribosomal, mitochondrial, and synaptic functions, whereas nuclear ambient RNA markers were mainly enriched for genes related to synaptic function **(Extended Figure 2A–B) (Supplementary Table 3)**. As also hypothesized in a previous study^22^, we then asked whether extra-nuclear ambient RNAs are enriched in genes comprising synaptosomes, which are also marked by ribosomal, mitochondrial and synaptic activity^23^. Using the top 500 markers in each ambient RNA group, we found that extra-nuclear ambient RNAs were more enriched than nuclear ambient RNAs in the transcripts of both vGLUT1-depleted (originating from postsynapse + soma) and vGLUT1-enriched (originating from presynapse) synaptosomes in Hafner et al.^23^ **(Extended Figure 2C)**. Since nuclear ambient RNAs also showed significant association and were enriched in overall synaptic function but not ribosomal and mitochondrial function **(Supplementary Table S3)**, we then hypothesized that nuclear ambient RNAs can instead be mainly derived from highly expressed genes captured in neuronal nuclei. Indeed, nuclear ambient RNAs largely overlapped with the highly expressed genes in neurons **(Extended Figure 2D)**. We note that this also explains significant enrichments between neuronal cell type markers and nuclear ambient RNA markers that we previously observed **(Figure 2E)**. Extra-nuclear ambient RNA and nuclear ambient RNA markers were also distinct, further underscoring that ambient RNAs are derived from different sources **(Extended Figure 2D).**

### Ambient RNA contamination of glial cells can be removed in silico

Given that neuronal genes are overrepresented in both ambient RNA types, we then hypothesized that ambient RNA contamination can make the transcriptomic profile of glial cell types more neuronal-like. As sorting for nuclei that do not express the neuronal marker NeuN (referred to as NeuN-) prior to droplet capture removes neuronal ambient RNAs, we utilized a NeuN-sorted snRNA-seq dataset for comparison^15^. We systematically found genes significantly overrepresented in three datasets (SD1, NSD1, NSD2) compared to the NeuN-sorted dataset in 6 major cell types: excitatory and inhibitory neurons, oligodendrocytes, OPCs (oligodendrocyte progenitor cells), astrocytes and microglia **(Supplementary Table 4)**. We called these genes “NeuN-depleted genes”. We then selected the top 500 NeuN-depleted genes and performed enrichment with ambient RNA markers. Interestingly, NeuN-depleted genes in all glia cell types were consistently enriched in nuclear ambient RNA markers across the three studies **(Figure 3A)**. We did not observe the same pattern with neuronal cell types. Extra-nuclear ambient RNAs were also enriched among NeuN-depleted genes in glia, although not consistently across datasets **(Figure 3B)**. Together these results show that glia in cortical snRNA-seq are contaminated with neuronal ambient RNAs.

**Figure 3.**
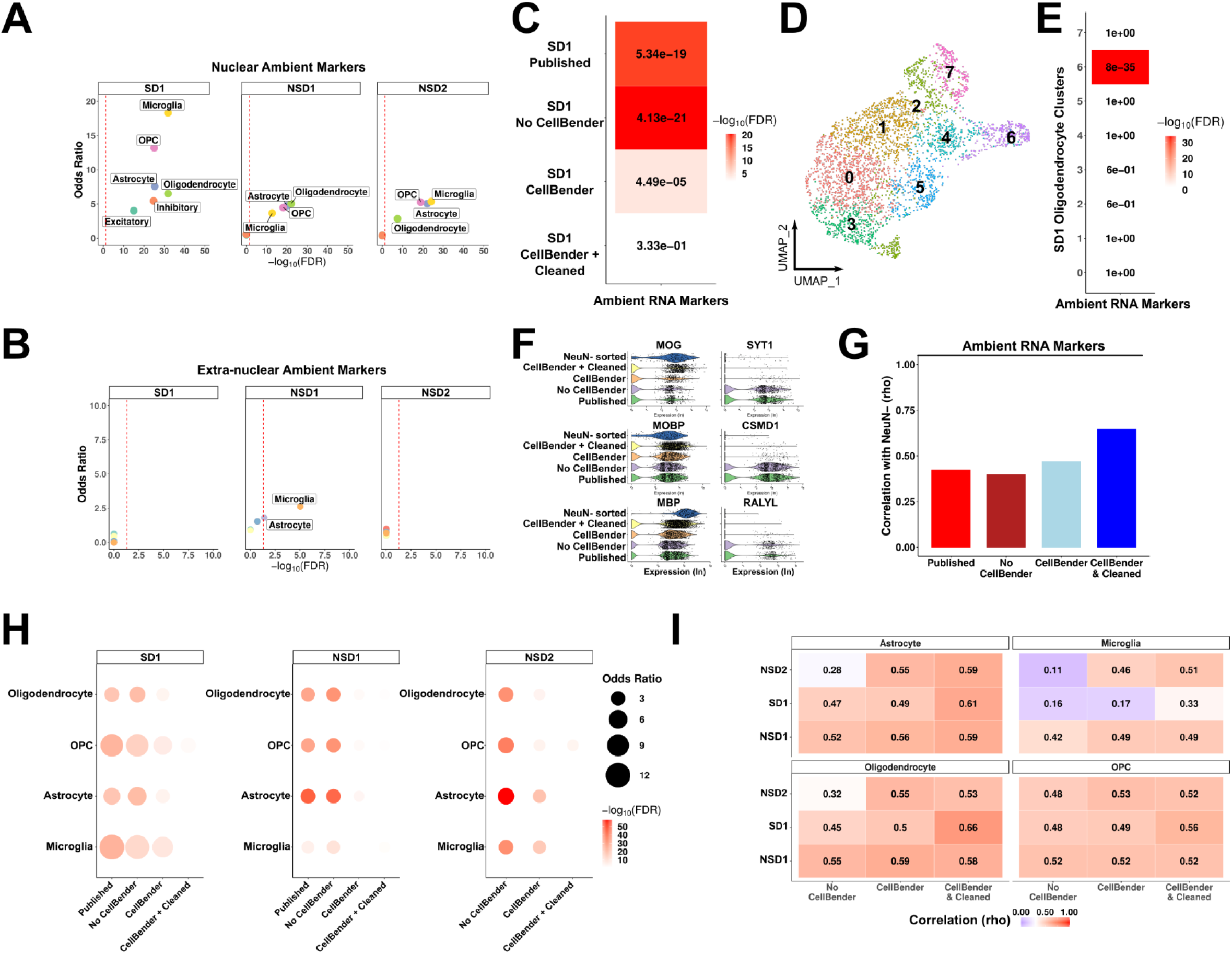
Ambient RNAs contaminate glia expression profiles. **(A-B)** Enrichment between genes depleted in NeuN-sorted dataset and nuclear ambient RNA markers **(A)** and extra-nuclear ambient RNA markers **(B)** across datasets per glial cell type (Fisher’s exact test, dashed line: FDR = 0.05). **(C)** Same enrichment as in **(A-B)** after each analysis (in y-axis as rows) performed in oligodendrocytes from SD1. dataset. Numbers show FDR value, colors are scaled based on −log_10_(FDR). **(D)** UMAP plot of SD1 oligodendrocytes after CellBender. **(E)** Enrichment between oligodendrocyte cluster markers and ambient RNA markers. **(F)** Violin plots of gene expression (log transformed) in oligodendrocytes after each analysis. Left column: oligodendrocyte markers, right column: ambient RNA markers. **(G)** Spearman rank correlations of all genes between SD1 oligodendrocytes and NeuN-sorted oligodendrocytes after each analysis (x-axis). **(H)** Same enrichment as in **(C)**, performed in all datasets and glial cell types after each analysis. **(I)** Spearman rank correlations of all genes with NeuN-sorted dataset. Correlations were performed per cell type per dataset (y-axis) after each analysis (x-axis). Both numbers and heatmaps show correlation magnitude.

Ambient RNA contamination within droplets that contain real cells is a general problem in single-cell RNA-seq experiments and various tools exist to remove ambient RNA contamination^7–9^. We thus utilized CellBender^9^ to remove the neuronal ambient RNA contamination. Focusing on oligodendrocytes (from SD1), we found that CellBender substantially reduced ambient RNA contamination **(Figure 3C)**. However, enrichment was still significant, indicating that ambient RNA contamination was not fully removed. To investigate this, we subclustered oligodendrocytes and found that markers of a small subcluster were highly enriched in ambient RNAs **(Figure 3D–E)**. Removing this subcluster fully removed detectable ambient RNA contamination from oligodendrocytes **(Figure 3C, F)** and increased correlation with NeuN-sorted oligodendrocytes **(Figure 3G)**. We then applied this procedure to each glial cell type per dataset and found that there was little to no contamination after CellBender and additional subcluster cleaning **(Figure 3H)**. Removed subclusters had slightly but consistently less intronic read ratio in datasets that did not perform nuclei-sorting, in line with the expectation that ambient RNA contamination contains extra-nuclear reads unless nuclei are physically sorted **(Extended Figure 3A)**. Indeed, intronic read ratios were similar between removed subtype and others in the nuclei sorted dataset **(Extended Figure 3B)**.

To test the effect of ambient RNA removal on all genes, we also assessed correlation of all expressed genes between the given dataset and NeuN-sorted dataset. We found that ambient RNA removal consistently increased overall correlations, indicating that ambient RNA removal results in better reproducibility between datasets **(Figure 3I)**. Neurons were also similarly reproducible compared to the NeuN+ sorted dataset before and after ambient RNA removal **(Extended Figure 3C)**. These results indicate that, ambient RNA contamination in glia can be effectively removed with CellBender and subcluster cleaning without undesired effects on the overall gene expression profile.

### Ambient RNA contamination is also detected in mouse brain snRNA-seq dataset

To assess if ambient RNA contamination is similar in mouse cortical snRNA-seq, we generated a snRNA-seq dataset from P56 (postnatal day 56) mouse frontal cortex. Similar to human datasets, the intronic read ratio was less in cell barcodes with low UMI **(Extended Figure 4A)** and ambient clusters were distributed bimodally with Low-Intron-CB and High-Intron-CB **(Extended Figure 4B–C)**. Low-Intron-CB markers were also enriched in extra-nuclear ambient RNAs whereas High-Intron-CB markers were enriched in nuclear ambient RNAs **(Extended Figure 4D)**. We similarly ran CellBender and performed subcluster cleaning on glial cell types. Both steps selectively removed ambient RNA signature from glial cell types **(Extended Figure 4E)**. Thus, we conclude that ambient RNA types and contamination of glial cell types by neuronal ambient RNAs are not specific to human datasets.

### Previous annotations of immature oligodendrocytes are glia contaminated with ambient RNAs

Glial cells can express genes that are typically associated with neuronal function, which raises the possibility that ambient RNA contamination might have been implicated with real biological function in previous snRNA-seq studies. Indeed, OPCs (oligodendrocyte progenitor cells) are known to make synapse-like contacts with axons and express glutamatergic receptors that binds to neurotransmitters secreted by neurons, affecting oligodendrocyte maturation in vitro^24–26^. We note that glutamatergic receptors functionally studied in oligodendrocyte maturation (e.g. *GRIA2*, *GRIA4*, *GRM5* in OPCs^24,25,27^) remain present in our analysis after the ambient RNA removal procedure **(Extended Figure 5A)**. Previous snRNA-seq studies identified “immature oligodendrocytes” which were marked by higher expression of many neuronal genes that were not functionally validated^20^. Based on our findings, we considered the alternative possibility that this excessive neuronal gene expression signature might be ambient RNA contamination. In line with this interpretation, we found that ~80% (46 out of 58) of immature oligodendrocyte markers overlapped with the top 200 most abundant ambient RNA markers **(Figure 4A)**. We then hypothesized that ambient RNA contaminated cells can be detected as transitioning cells between any two glial cell types in pseudotime analysis as all glial cells are vulnerable to ambient RNA contamination. Using the gene-cell matrix from the original publication, we were able to reconstruct the lineage trajectory between OPC-OL **(Figure 4B)**. Cell barcodes between OPC and OL (‘transitioning cells’) showed high enrichment in immature oligodendrocyte and ambient RNA markers **(Figure 4B–C)** and these cell barcodes were removed during our subcluster cleaning procedure **(Figure 4D)**. These cell barcodes displayed similar UMI counts compared to OPC and OL indicating that they are also not glia-neuron doublets **(Figure 4E)**. We then created pseudotime analysis between OPC-AST (astrocytes) and found similar ‘transitioning cells’ **(Extended Figure 5B-D)**. Since OPCs can achieve multipotency under certain conditions^28–30^, we could not exclude the possibility that this could be a real biological function (i.e. OPCs differentiating into astrocytes). For a definitive answer, we then tested two non-oligodendrocyte lineage cell types, AST and MIC (microglia), which also revealed similar ‘transitioning cells’ that were highly enriched in both ambient RNA and immature oligodendrocyte markers and were effectively removed by our ambient RNA removal process **(Extended Figure 5E-G)**. Given that immature oligodendrocytes also lack known markers of committed OPCs (COPs) or premyelinating oligodendrocytes (Pre-OL) (e.g *BCAS1*, *ENPP6*, *GPR17*^31^) **(Extended Figure 5H)**, our results indicate that cells previously annotated as immature oligodendrocytes are not transitioning cells but rather glia with high contamination of neuronal ambient RNA.

**Figure 4.**
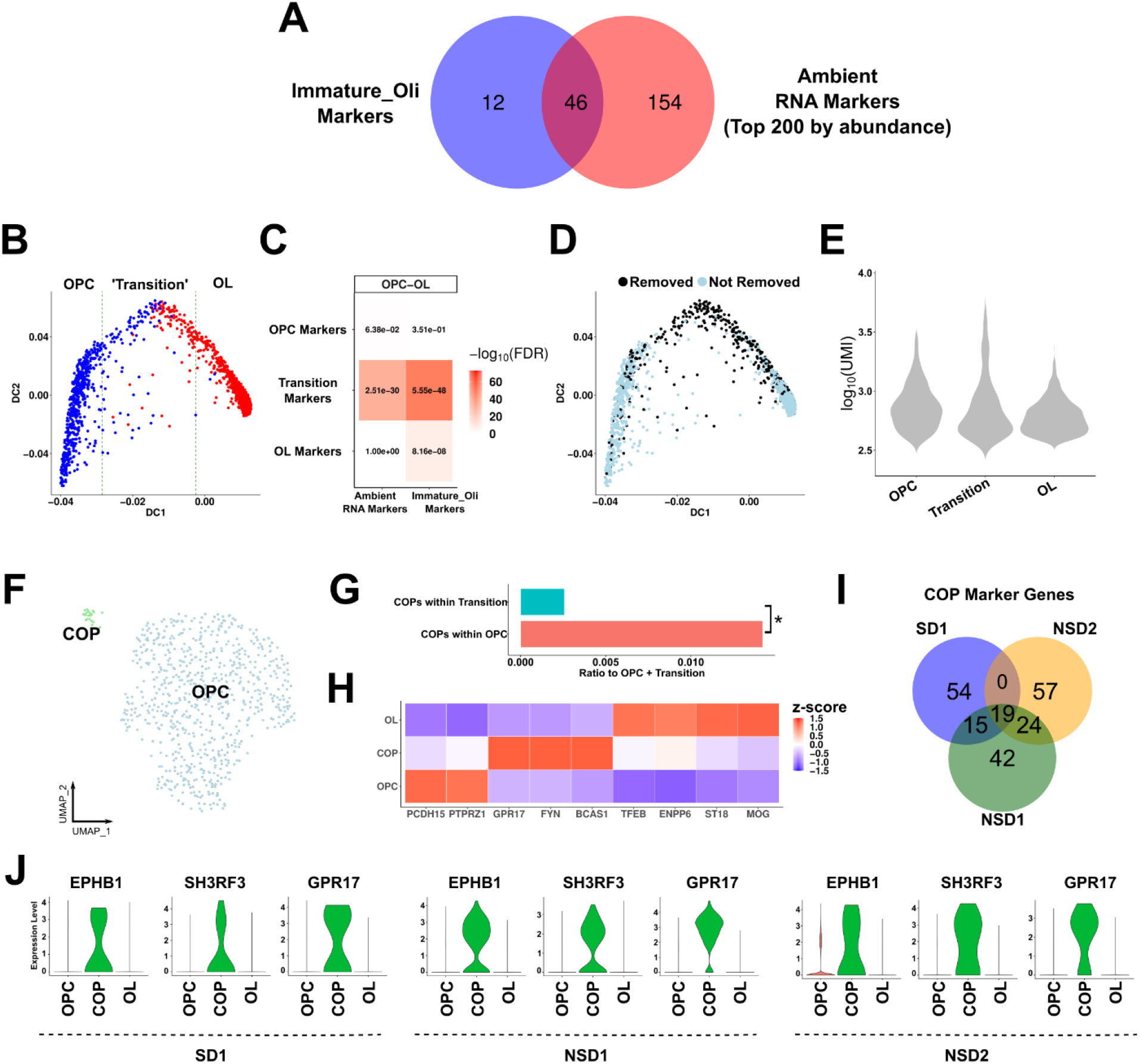
Committed progenitor cells are detectable in adult human brain. **(A)** Overlap of immature oligodendrocyte markers in SD1 and top 200 most abundant ambient RNA markers. **(B)** Oligodendrocyte lineage trajectory as reconstructed with *destiny*. ‘Transition’ zone was determined as 400 cells around the middle cell based on DC1. **(C)** Enrichments between trajectory zones (OPC, Transition, OL) and ambient RNA or immature oligodendrocyte markers (Fisher’s exact test, numbers indicate FDR, color scale is −log_10_(FDR)). **(D)** Same lineage trajectory as **(B)** with cells removed after subcluster cleaning highlighted. **(E)** UMI counts of cell barcodes within OPC, Transition or OL zones. **(F)** UMAP of OPC subclustering. COP: committed OPCs. **(G)** Ratio of COPs within OPCs or within transitioning cells to total number of OPCs and transitioning cells. Asterisk indicates significance (p-value < 0.05, Chi-square test). **(H)** Heatmap of oligodendrocyte lineage markers (z-scored across cell types per marker gene). **(I)** Overlap of COP markers (compared to OPCs) across datasets. Only top 100 markers were selected (FDR < 0.05). **(J)** Expression levels of top COP markers in three datasets.

### Ambient RNA removal reveals rare cell type in adult human brain snRNA-seq datasets

Initial single cell studies on the adolescent mouse oligodendrocyte lineage identified COPs and NFOLs (newly-formed oligodendrocytes) as transitioning OL cells^16^. This work established marker genes including *Gpr17,* which peaked in COPs, was reduced in NFOLs and was absent in mature OLs^16^. A study in human induced pluripotent stem cell derived OPC culture also showed that GPR17 regulated OL maturation in human cells^32^. However, few single-cell RNA-seq studies in adult human brain identified these populations^13,33^. Since these studies used different annotation labels and marker genes, it is also unclear whether transitioning oligodendrocytes are consistent across human datasets. Robust annotation of these cells in human datasets is crucial to understand the role of the oligodendrocyte lineage in neurological diseases^11,13,34,35^. As we have shown, ambient RNA contamination can potentially mislead the results and should be cleared to obtain biologically relevant results.

To observe whether we can identify COPs after ambient RNA removal, we subclustered OPCs. *GPR17*+ COPs were detectable and clustered separately than OPCs in all adult datasets **(Figure 4F, Extended Figure 6A,C)**. Importantly, COPs were significantly depleted within the transition zone further indicating that previous pseudotime analysis was driven by ambient RNA contamination and it is not relevant for oligodendrocyte maturation **(Figure 4G)**. To validate the identity of COPs, we then plotted genes known to be associated with COPs or Pre-OLs (*BCAS1^36^*, *GPR17^37^*, *FYN^38^*) as well as genes that are upregulated in Pre-OLs but are also expressed in OLs (*TFEB^39^*, *ENPP6^40^*). We found that *BCAS1*, *GPR17* and *FYN* selectively marked COPs and *TFEB* and *ENPP6* were upregulated in COPs compared to OPCs consistently across datasets **(Figure 4H, Extended Figure 6B,D)**. Overall, 58 out of 211 (27%) of the top markers of COPs were shared across at least two datasets **(Figure 4I)**. To highlight novel markers for COPs in adult human brain, we then found the most specific COP markers compared to both OPCs and OLs which – in addition to *GPR17*, *BCAS1*, *FYN* – revealed *TNS3^16^*, *SH3RF3*, *EPHB1*, *CRB1*, *SIRT2* and *ARHGAP5* adding to the list of potential markers which can be valuable for future studies to understand oligodendrocyte biology in human brain **(Figure 4J, Supplementary Table 5)**. Given that we could detect similar COP populations in all datasets, we also re-analyzed a previous study that identified COPs in adult human brain white matter^13^. Surprisingly, COP markers were not highly expressed in the original annotation of COPs **(Extended Figure 7A)**. Clustering OPCs and COPs revealed a subpopulation of cells that were very similar to COPs in other human datasets by their marker gene expression levels (‘COPs-New’) **(Extended Figure 7B–C)**. To assess whether previously annotated COPs (‘COPs-Old’) can be ambient RNA contamination, we also checked expression levels of neuronal genes. This revealed high expression of both ambient and non-ambient neuronal genes, indicating ‘COPs_Old’ might be OL-neuron doublets rather than ambient RNA contamination **(Extended Figure 7C)**. Indeed, COPs-Old displayed similar UMI levels to neuronal cell types, in contrast to ambient RNA driven clusters which contained lower UMI levels **(Extended Figure 7D, Figure 4D, Figure 1A)**. These results provide further evidence of extreme rarity of COPs in human brain datasets which can be masked by technical artifacts.

### Stepwise guideline for detection and removal of ambient RNAs

Our results show that a combination of existing tools and careful analysis can remove ambient RNA contamination and improve the biological relevance of results. To illustrate our approach in a more direct way, we present a stepwise guideline that outlines the major steps important in our analysis **(Extended Figure 8)**. While ambient RNA removal tools aim to be a one-step solution for this problem, we advise researchers to identify ambient RNA populations and their markers genes in their own dataset which is achievable using common methods **(Extended Figure 8, Steps 1-4)**. This can then be used to assess whether a specific cell population is marked by high ambient RNA contamination which may not have been removed or cleaned of ambient RNAs by the specialized tools (e.g. CellBender, **Extended Figure 8, Steps 5-9**). We note that our experience is limited to single-nuclei datasets in brain, however most analysis steps as well as existing tools can be used for any single-cell dataset.

Taken together, we show pervasive contamination of glia by neuronal ambient RNAs, successfully remove them using available methods which reveals underappreciated biology of transitioning oligodendrocytes in adult human brain. We also provide a stepwise guideline outlining our integrated approach to tackle ambient RNA contamination in single-nuclei datasets from brain tissue.

## Discussion

In this study we provide in depth examination of ambient RNAs specifically in brain snRNA-seq datasets. We identify nuclear and extra-nuclear ambient RNAs that have different gene signatures **(Extended Figure 2)**. We find that previously annotated neuronal cell types have high contamination of ambient RNAs^12,14^ **(Figure 1–2)**. We then show that the high prevalence of neuronal reads in ambient RNAs contaminate glia which can be effectively removed by CellBender and additional subcluster cleaning **(Figure 3)**. These results are not unique to human brain and are reproduced in mouse snRNA-seq data **(Extended Figure 4)**. We also show that previous identification of immature oligodendrocytes is a result of neuronal ambient RNA contamination **(Figure 4)**. After ambient RNA removal, careful examination of OPCs reveals COPs in all human brain snRNA-seq datasets **(Figure 5)**. We finally highlight both known and novel markers of COPs within the oligodendrocyte lineage that can be further studied and provide a stepwise guideline of ambient RNA marker identification and removal.

An important observation in our findings is that single-nuclei isolation does not entirely remove extra-nuclear reads. We leveraged this information to reveal that previously annotated cell clusters (Neu-NRGNs) contained high levels of extra-nuclear ambient RNA contamination **(Figure 1C)**. However, extra-nuclear contamination measured by intronic read ratio is not sufficient as a single metric to identify undesirable cell barcodes. For example, a recent study highlighted the distinction of damaged cells and empty droplets that only contain ambient RNA^5^. The authors noted that damaged cells contain similar intronic read ratio (i.e. nuclear fraction) to real cells, but they display lower UMI counts compared to other cells with similar annotation^5^. Similarly, we find that Neu-mat cluster has lower UMI compared to other neurons despite having similar intronic read ratio, indicating that they are likely damaged nuclei **(Figure 1A, C)**. Additionally, we observe that while empty droplets have lower intronic read ratio, this ratio is still substantially higher than zero, indicating that nuclear reads also contribute to ambient RNAs **(Figure 1C, 2A)**. Moreover, in glial cell barcodes that contain both high ambient RNA contamination and a real nucleus, intronic read ratio is only slightly less than other nuclei **(Extended Figure 3A)**.

This is likely due to the lower abundance of extra-nuclear ambient RNAs compared to the entire transcriptome of the real nuclei. Finally, intronic read ratio is not an indicator of ambient RNA contamination in datasets that perform nuclei sorting by flow cytometry since this procedure removes extra-nuclear ambient RNAs **(Figure 2B)**. In contrast, ambient RNA markers can be identified in any dataset by analyzing empty droplets (i.e. ambient cell barcodes) and any group of cells can be assessed for enrichment of ambient RNA signature. We provide a stepwise guideline that outlines this process **(Extended Figure 8)**.

In addition to empty droplets, ambient RNAs can potentially contaminate all real cells/nuclei and several tools exist to overcome this problem^7–9^. Here we used CellBender^9^ which efficiently removed neuronal ambient RNA contamination from glial nuclei **(Figure 3)**. Upon examination, we realized that CellBender did not remove or clean some cell barcodes with high neuronal read contamination. This was detectable in most glial cell types and clustered separately than other cell barcodes which made it straightforward to identify and remove them **(Figure 3C–E)**. Importantly, a benchmark study also observed that CellBender retained more cell barcodes than other methods^4^. We therefore note that other tools more specialized for detection of real cells^2,4,5^ may yield better results and can be used in conjunction with ambient RNA contamination removal tools like CellBender.

In the datasets we assessed, ambient RNA markers were distinctly neuronal which resulted in clear detection of contamination in glia. This is likely because of transcriptomic abundance of neurons compared to other cell types in brain tissue. While we found high correspondence of ambient RNA markers between datasets, and even between species **(Figure 1–2, Extended Figure 1,4, Supplementary Table 5)**, we limited our analysis to cortical gray matter samples. We expect ambient RNA signature to change depending on the analyzed tissue which can result in a different contamination pattern based on the prominent cell type.

Lastly, we showed that immature oligodendrocytes can be explained by ambient RNA contamination and further analysis after ambient RNA removal revealed COPs in three independent datasets. We note that while COPs have been identified before^16,17^, many studies did not annotate them ^11,12,14,21^. This could be especially hindered by both ambient RNA contamination and the extreme rarity of COPs as a cell type in adult brain. Indeed, COPs are only ~0.04% of all cells in adult brain (NSD2 dataset, 30-80 years old) and ~0.3% of all cells in adolescence (NSD1^12^, 4-22 years old). We also showed that previously annotated COPs in adult human brain white matter are likely OL-neuron doublets and real COPs are detectable and similarly rare (~0.1% of all cells) **(Extended Figure 7)**. In line with this observation, live cell imaging in mouse showed that ~80% of transitioning oligodendrocytes rapidly undergo cell death, which is expected to result in a transient and rare cell population^41^. A carbon dating study of human genomic DNA also showed low levels of oligodendrogenesis in adulthood, which provides further support for the extreme rarity of transient cells in adult human brain^42^. Despite being rare, COPs are a critical cell type for future studies to examine since oligodendrocyte maturation is altered in both neurological diseases^35,43^ and human evolution^44,45^. Thus, we underscore the importance of ambient RNA removal, especially for accurate analysis of underrepresented cell types, and here provide novel markers for COPs in the adult human brain.

Another recent study also uncovered that glial cell types respond to enzymatic dissociation during single-cell and single-nucleus library preparation and confound the transcriptomic profile^46^. Here, we show that all glial cell types are also contaminated with neuronal ambient RNA transcripts. Together, these results indicate that both data generation and data analysis of glial cell types should be revisited and updated.

## Methods

### Preprocessing and count matrix generation

Datasets were downloaded from NCBI-GEO database **(Supplementary Table 1)**. Within the datasets, we only used the cells generated from cortical brain tissue. Specifically, NSD1 was prefrontal and anterior cingulate cortex, NSD2 was posterior cingulate cortex, SD1 was prefrontal and visual cortex, and SD2 was anterior cingulate cortex. Barcode correction and filtering was done using *umi_tools whitelist* (retained top 20,000-80,000 cell barcodes per sample depending on the dataset to keep ambient cell barcode population) and *umi_tools extract^47^*. Alignment was done using STAR aligner^48^ with the reference genomes of GRCh38 (for the human datasets) or GRCm38 (for the mouse dataset). *featureCount* was used to count reads mapping to gene body only for uniquely mapping reads^49^, and *umi_tools count* was used to create the count matrix. Count matrix using only intronic reads were similarly obtained using featureCount on a custom gtf that only contained introns (created using *construct_introns* from *gread* package in R^50^). Intronic read ratio was then calculated per cell barcode by taking the ratio of number of UMIs mapping to introns and number of UMIs mapping to gene body.

For ambient RNA cleanup, CellBender was used on the raw matrix of gene counts with default parameters^9^.

### Single-nuclei library preparation

We processed 4 c57BL/6J P56 mice (2 males and 2 females). Mice were rapidly decapitated and brains were quickly removed. The isolated brain was quickly transferred to an ice-cold coronal brain section mold (Braintree Scientific, BS-A 5000) and washed with ice-cold 1X PBS (Cytiva, SH30256.01). The boundary of the olfactory bulb and frontal cortex was aligned to the first indentation where the first razor blade (Fisher Scientific, 12-640) was inserted to remove olfactory bulb. The second razor blade was inserted into the third indentation. This coronal section matches with coronal numbers 22-36 in the Allen Brain Atlas: Mouse Reference Atlas, Version 2 (2011). We then removed the subcortical region from this section and separated the left and right hemisphere samples into different Eppendorf tubes. The tubes were flash frozen in liquid nitrogen.

The nuclei isolation procedure was modified from our previous work^51^. The frozen section from the left hemisphere was transferred to a Dounce homogenizer with 2ml of ice-cold Nuclei EZ lysis buffer (Sigma-Aldrich, NUC101). We then inserted pestle A for 23 strokes followed by pestle B for 23 strokes on ice. The homogenized sample was transferred to a 15ml conical tube. We added 2ml of ice-cold Nuclei EZ lysis buffer and incubated on ice for 5min. Nuclei were collected by centrifuging at 500 g for 5min at 4C. We discarded the supernatant and added 4ml of ice-cold Nuclei EZ lysis buffer to resuspend nuclei. We then repeated the incubation and centrifuge steps and resuspended the nuclei in 200ul of nuclei suspension buffer: 1X PBS, 1% BSA (ThermoFisher, AM2618), and 0.2 U/ul RNAse inhibitor (ThermoFisher, AM2696). Finally, the nuclei suspension was filtered twice through Flowmi Cell Strainers (Bel-Art, H13680-0040).

We mixed 10ul of nuclei suspension with 10ul of 0.4% Trypan Blue (Gibco, 15-250-061) and loaded on a hemocytometer (SKC, DHC-N015) to determine the concentration. 10,000 nuclei/sample were used to prepare snRNA-seq libraries using 10X Genomics Single Cell 3′ Reagent Kits v3^52^. Libraries were sequenced by the McDermott Sequencing Core at UT Southwestern on a NovaSeq 6000.

### Ambient cluster analysis

To retain cell barcodes that predominantly contain ambient RNAs, we kept two times more cell barcodes than the original publication per sample. For datasets generated in this study, we retained two times more cell barcodes than the number of nuclei targeted. Since not all newly retained cell barcodes are predominantly ambient RNAs (e.g. they could be doublets, or low quality nuclei of various cell types), we then performed clustering analysis to identify ambient RNA clusters that clustered distinctly compared to annotated cell types per dataset. The following methods from Seurat v3^53^ were used to perform and visualize clustering: normalization (*SCTransform*), dimensionality reduction (*RunPCA*), batch correction (*RunHarmony*, default parameters), k-nearest neighbors (*FindNeighbors)* on batch corrected dimensions and clusters identification by shared nearest neighbors (*FindClusters*). UMAP embedding was then computed for visualization in 2D space (*RunUMAP*). Clusters that contained >75% newly retained (i.e. filtered in the original publication) were annotated as ambient clusters. We note that 75% is unusually high since only 50% of all cell barcodes is newly retained and cell types are expected to be mostly retained in the original publication.

To identify ambient cluster marker genes, we ran DGE (differential gene expression) analysis using pseudobulk edgeR^54^. Briefly, counts were aggregated per sample and pseudobulk DGE was run with *pseudoBulkDGE* function *(method = ‘edgeR’)* in *scran* package^55^. Ambient cluster markers were identified with logFC > 0.3 and FDR < 0.05 cutoffs.

Enrichment of ambient cluster markers with annotated cell types were done using Fisher’s exact test from *GeneOverlap* package^56^. The total number of expressed genes were used as background (we followed this strategy in all Fisher’s exact test enrichments).

### Comparison with NeuN-dataset

To find differential genes in each dataset compared to the NeuN-sorted dataset from Hodge et al.^15^, we first identified 6 major cell types in all datasets: excitatory neurons, inhibitory neurons, oligodendrocytes, OPCs, microglia and astrocytes. Using pseudobulk DGE (see above) in matched cell types, we then identified differential genes with significantly higher amount of reads than in the NeuN-sorted dataset. This was performed separately for each dataset.

For enrichment with ambient cluster markers, we selected the top 500 differential genes (ranked by logFC) among NeuN-depleted genes in each comparison (logFC > 1 and FDR < 0.05). Similarly, the top 500 ambient RNA markers were selected from both nuclear ambient RNA and extra-nuclear ambient RNA markers. Enrichment was performed as above.

### Subcluster cleaning of glia after CellBender

To subcluster glia after CellBender, we used the annotation provided in the original publication and processed each glia cell type separately per study. For the datasets generated in this study, we performed clustering as described and annotated glia based on the established markers genes (e.g. *MBP*, *PCDH15*, *APBB1IP*, *SLC1A3*). Clustering was done similarly as above and marker genes of subclusters were tested for enrichment in ambient RNA markers using Fisher’s exact test. For this, we selected top 500 (by logFC) ambient RNA markers from both nuclear and extra-nuclear ambient RNA and combined them. We removed the subclusters with distinctly high level of enrichment in ambient RNAs (FDR < 0.001 and odds ratio > 3) compared to other subclusters. All steps of ambient RNA contamination removal are outlined in **Extended Figure 8**.

### Assessment of ambient RNA contamination signatures in glia

To compare ambient RNA contamination in glia after different analyses, we first found cell barcodes common between annotated cell barcodes in the original publication and retained cell barcodes after CellBender. We then only retained common cell barcodes in all downstream analyses that compared the original dataset and analyses that included ambient RNA contamination removal **(Figure 3)**. This was to ensure that only gene expression levels were different and the enrichments are not driven by cell barcode differences between analyses. However, we note that the analysis with CellBender + subcluster cleaning contained fewer cell barcodes as ambient RNA rich subclusters were removed after CellBender. To test ambient RNA marker enrichment, we first found differential genes with significantly higher amount of reads than NeuN-sorted dataset per cell type per dataset (logFC > 1 and FDR < 0.05). These gene lists were then tested for enrichment in ambient RNA markers (combined top 500 genes was used as before) using Fisher’s exact test.

To test whether the global gene expression profile is altered after these different analysis methods, we found genes expressed in at least 5% of cells per cell type per dataset to remove lowly expressed genes. We then kept the genes that survive this threshold in all datasets and performed Spearman rank correlation between each dataset and the NeuN-sorted dataset using the average log expression of genes in the normalized matrix.

### Pseudotime analysis

To be consistent with SD1^20^, we used *DiffusionMap* function from *destiny^57^* only on visual cortex samples to build pseudotime between OPC-OL or between other pairs of glial cell types using the matrix provided by the authors. Diffusion maps were created with parameters *n_pcs=100* and *k=100*. The first two eigenvectors of diffusion maps were plotted for visualization. To identify markers of ‘transitioning’ cell barcodes, we found the middle cell barcode based on the first eigenvector (DM1) and labeled 200 cell barcodes around the middle cell barcode as ‘transitioning cells’. The remaining two groups of cell barcodes were labeled by their original annotation label (e.g. OPC). We then found marker genes for each of these pseudotime groups (FDR < 0.05 and logFC > 0.25 using *FindMarkers* in *Seurat^53^*) and ran enrichment with ambient RNA markers and immature oligodendrocyte markers identified in SD1 using Fisher’s exact test.

### OPC subcluster analysis

To identify potential transitioning OPCs, we separately subclustered OPCs in three different datasets (SD1^20^, NSD1^12^ and NSD2) after CellBender and subcluster cleaning based on high ambient RNA contamination. We further removed subclusters with high expression of markers from two distinct major cell types as potential doublets. Committed oligodendrocyte progenitors (COPs) were identified by high expression of *GPR17* (as previously established^16^) among other markers (e.g *BCAS1*, *FYN*).

To identify subclusters of Jäkel et al., we performed dimensionality reduction and clustering on cells with the annotation of ‘OPCs’ and ‘COPs’ using Seurat v3 as described above. We then identified ‘COPs-New’ by the established marker genes (*BCAS1, FYN, GPR17*). For the heatmaps, all mature oligodendrocytes were combined and annotated as ‘OL’. Normalized and log transformed expression levels for each gene was then z-transformed across 4 cell type annotations (OPC, COP-New, COP-Old, OL). For the UMI plot, we retained the original labels for neuronal cell types. Both control and multiple sclerosis samples were used and no additional cell filtering (other than subsetting by annotation) was applied for all analyses.

### Identification of COP Marker Genes

Genes upregulated in COPs compared to OPCs were identified using the FindMarkers function from Seurat. Significant genes (FDR < 0.05 and expressed in >10% of COPs) were ranked by their avg_logFC and top 100 genes per dataset were selected.

To highlight genes specific to COPs compared to OPCs and OLs, we found the percentage of cells that contained at least one copy for each significant gene. We then computed the difference of percentages between both COPs-OPCs and COPs-OLs. We then took the intersection of top 20 genes with the highest difference in favor of COPs in both comparisons. This was repeated for all three datasets. Genes that marked COPs in at least two datasets were reported as COP markers within the oligodendrocyte lineage in human brain **(Supplementary Table 5)**

### Other enrichments

Gene ontology (GO) enrichment of ambient RNA signatures was done using clusterProfiler package in R^58^ with all expressed genes used as the background. Full table is available in **Supplementary Table S3**.

To test enrichment of ambient RNA markers with vGLUT1-Depleted and vGLUT1-Enriched genes from Hafner et al.^23^, we first converted the mouse gene symbols to human gene symbols using SynGO^59^. Fisher’s exact test was performed as before.

To overlap ambient RNAs with highly represented genes in neurons in snRNA-seq, we identified the top-represented genes among all neurons by taking the mean of each gene across all cell barcodes annotated as neurons separately in both SD1 and NSD1 (except for Neu-NRGNs and Neu-mat). The intersection of the top 500 genes in both datasets (403 genes) was used to overlap with ambient RNA markers.

## Acknowledgments

The authors thank Dr. Tao Wang, Dr. Lu Sun, Dr. Dmitry Velmeshev, Dr. Arnold Kriegstien for their critical comments on the manuscript. G.K. is a Jon Heighten Scholar in Autism Research and Townsend Distinguished Chair in Research on Autism Spectrum Disorders at UT Southwestern. E.C. is a Neural Scientist Training Program Fellow in the Peter O’Donnell Brain Institute at UT Southwestern. This work was partially supported by the James S. McDonnell Foundation 21^st^ Century Science Initiative in Understanding Human Cognition Scholar Award, NHGRI (HG011641), NINDS (NS115821) and NIMH (MH126481, MH103517) to G.K. and an American Heart Association Postdoctoral Fellowship (915654) to Y.L.

## Author contributions

E.C. and G.K. conceptualized the study. Y.L. collected snRNA-seq data. E.C. performed all analyses. Y.L. edited the manuscript. E.C. and G.K. wrote the manuscript.

---

**Extended Figure 1.**
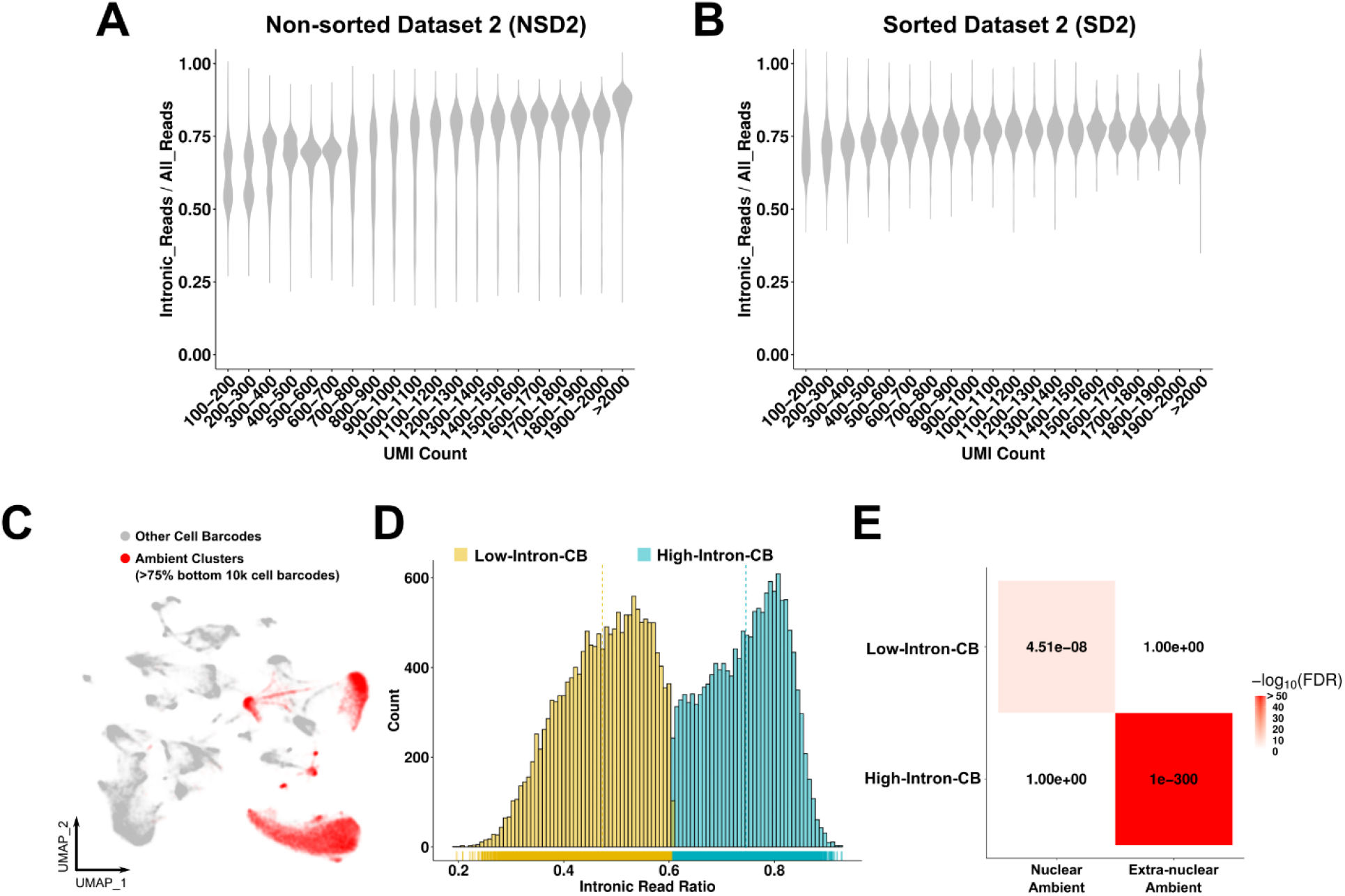
Independent confirmation of nuclear and extra-nuclear ambient RNA types. **(A-B)** Intronic read ratio across increasing UMI count in **(A)** non-sorted dataset 2 (NSD2) and **(B)** sorted dataset 2 (SD2). UMIs are divided into intervals of 100 from 100-2000. **(C)** Clusters that are >75% composed of filtered cell barcodes are highlighted and named ambient clusters (dataset: NSD2). **(D)**. Distribution of intronic read ratio within ambient clusters. Yellow: Low-Intron-CB (CB: Cell Barcodes), blue: High-Intron-CB. **(E)** Enrichments between ambient RNA types. Nuclear ambient RNA and extra-nuclear ambient RNAs are identified from SD1 and NSD1 (see Figures 1–2).

**Extended Figure 2.**
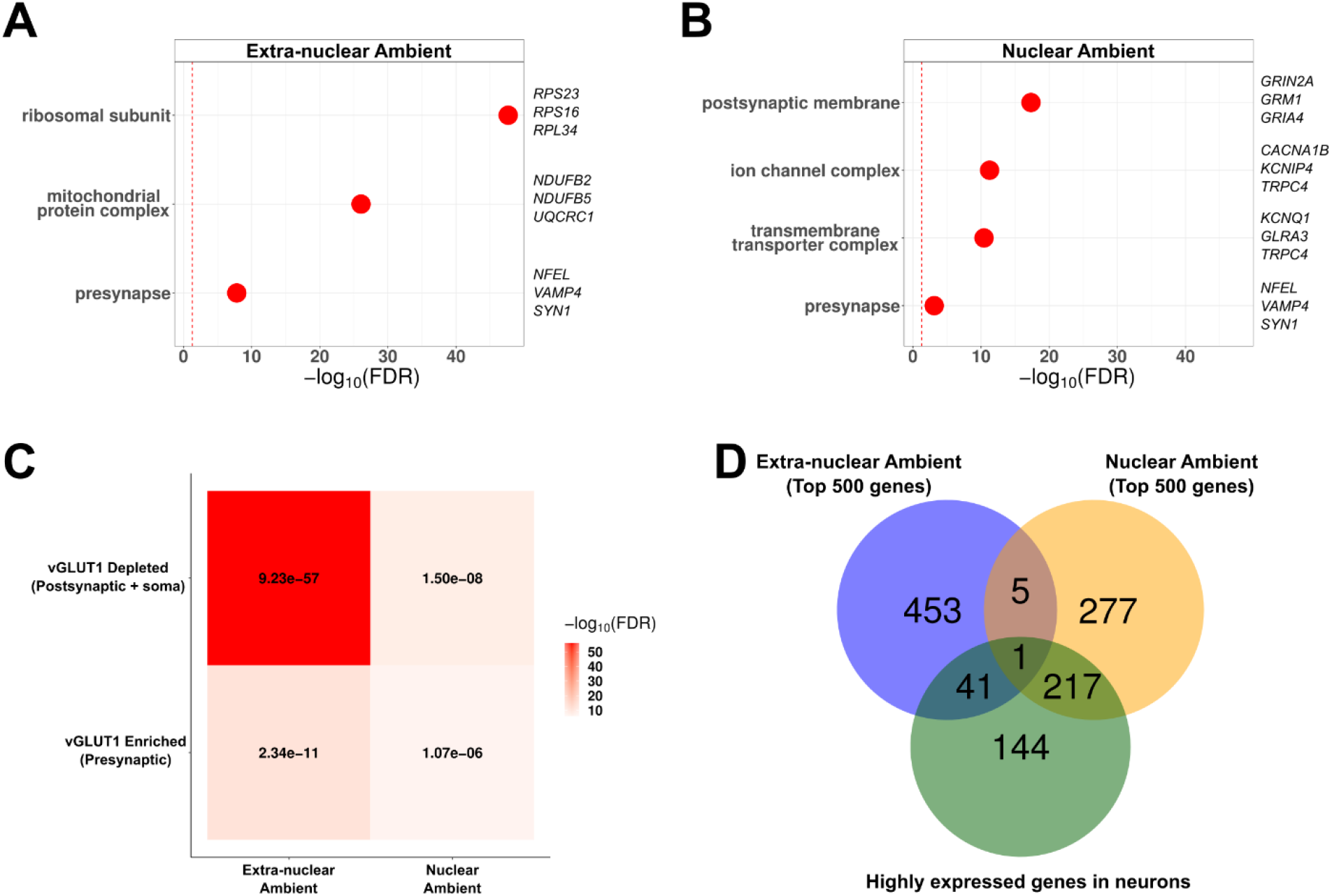
Characterization of ambient RNA markers. **(A-B)** Gene ontology (GO) enrichments for extra-nuclear ambient RNA markers **(A)** and nuclear ambient RNA markers **(B)**. Example genes per GO term are shown on the right. **(C)** Enrichments between presynaptic synaptosomes (vGLUT1 Enriched) and others (vGLUT1 Depleted) (Fisher’s exact test, numbers show FDR, heatmap color indicates −log_10_FDR). Genes from synaptosome dataset were converted from mouse to human symbols prior to enrichment. **(D)** Overlap between the top 500 most distinct ambient RNA markers and the top 500 highly expressed genes in neurons.

**Extended Figure 3.**
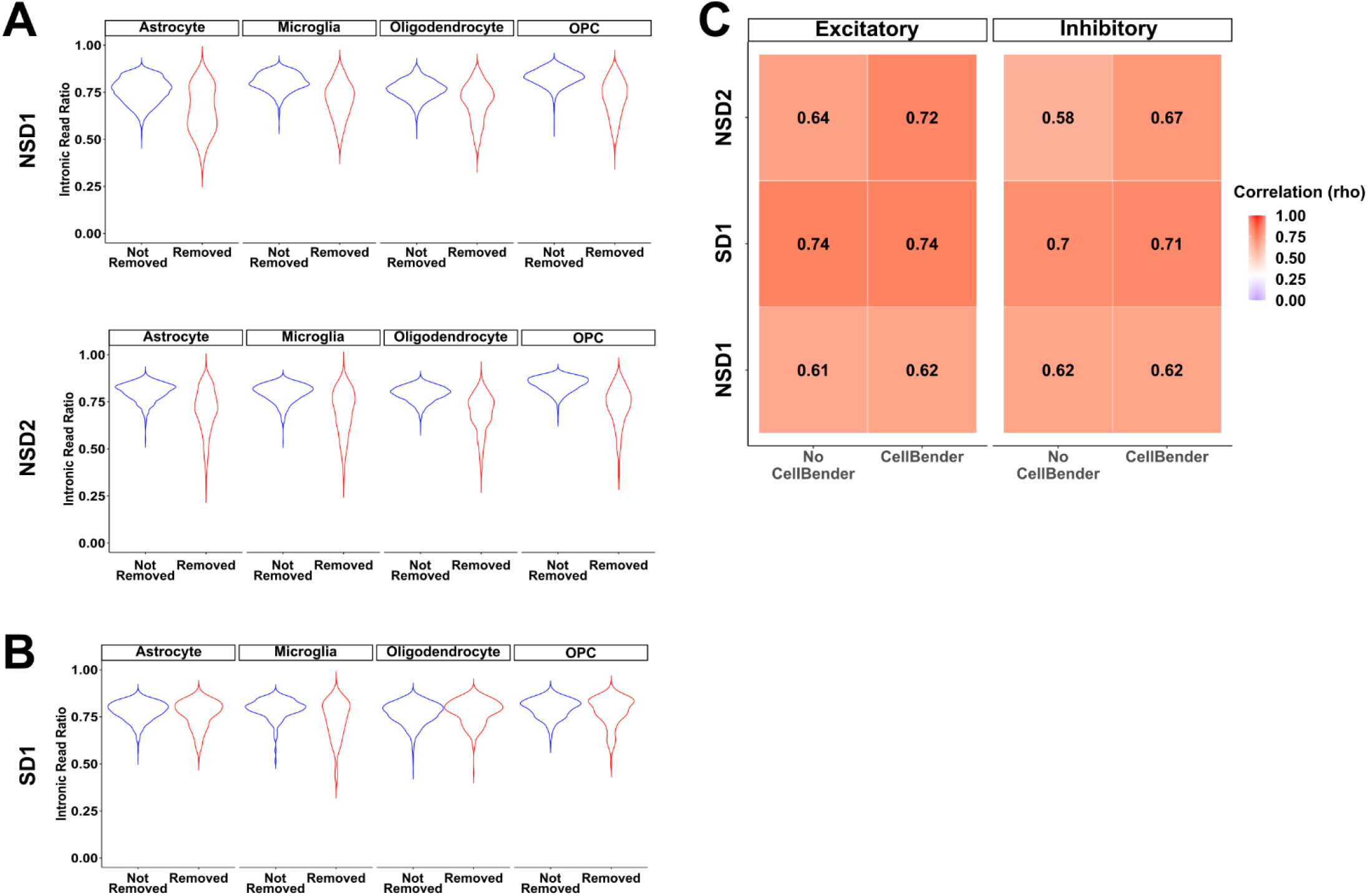
Supplementary analyses on CellBender and subcluster cleaning. **(A)** Intronic read ratio per cell barcode in subclusters that were removed due to ambient RNA contamination (red) or not removed (blue) per glial cell type in datasets that did not perform nuclei sorting. **(B)** Same as in **(A)** but in the dataset that performed nuclei sorting. **(C)** Spearman rank correlations of all genes with NeuN+ sorted dataset. Correlations were performed per cell type per dataset (y-axis) after each analysis (x-axis). Both numbers and heatmaps show correlation magnitude.

**Extended Figure 4.**
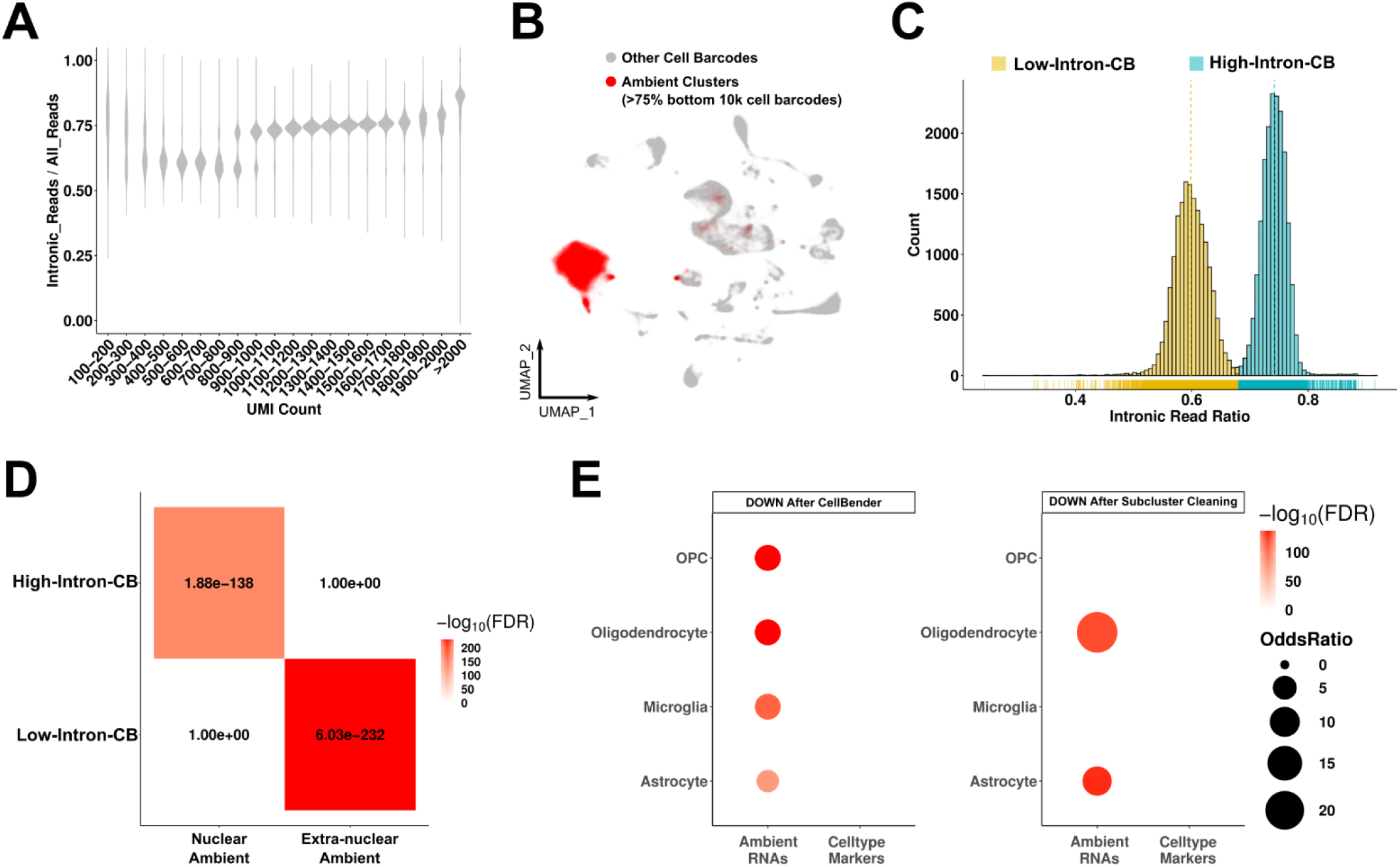
Ambient RNAs in mouse brain snRNA-seq dataset. **(A)** Intronic read ratio across increasing UMI count in mouse brain snRNA-seq dataset (no nuclei-sorting). UMIs are divided into intervals of 100 from 100-2000. **(B)** Clusters that are >75% composed of filtered cell barcodes are highlighted and named ambient clusters. **(C)**. Distribution of intronic read ratio within ambient clusters. Yellow: Low-Intron-CB (CB: Cell Barcodes), blue: High-Intron-CB. **(D)** Enrichments between ambient RNA types. Nuclear ambient RNA and extra-nuclear ambient RNAs are identified from SD1 and NSD1 (see Figures 1–2). **(E)** Enrichments between genes with significantly lower expression (DOWN) after CellBender (left) or subclustering steps (right) and ambient RNA markers or cell type markers. Cell types are shown in y-axis.

**Extended Figure 5.**
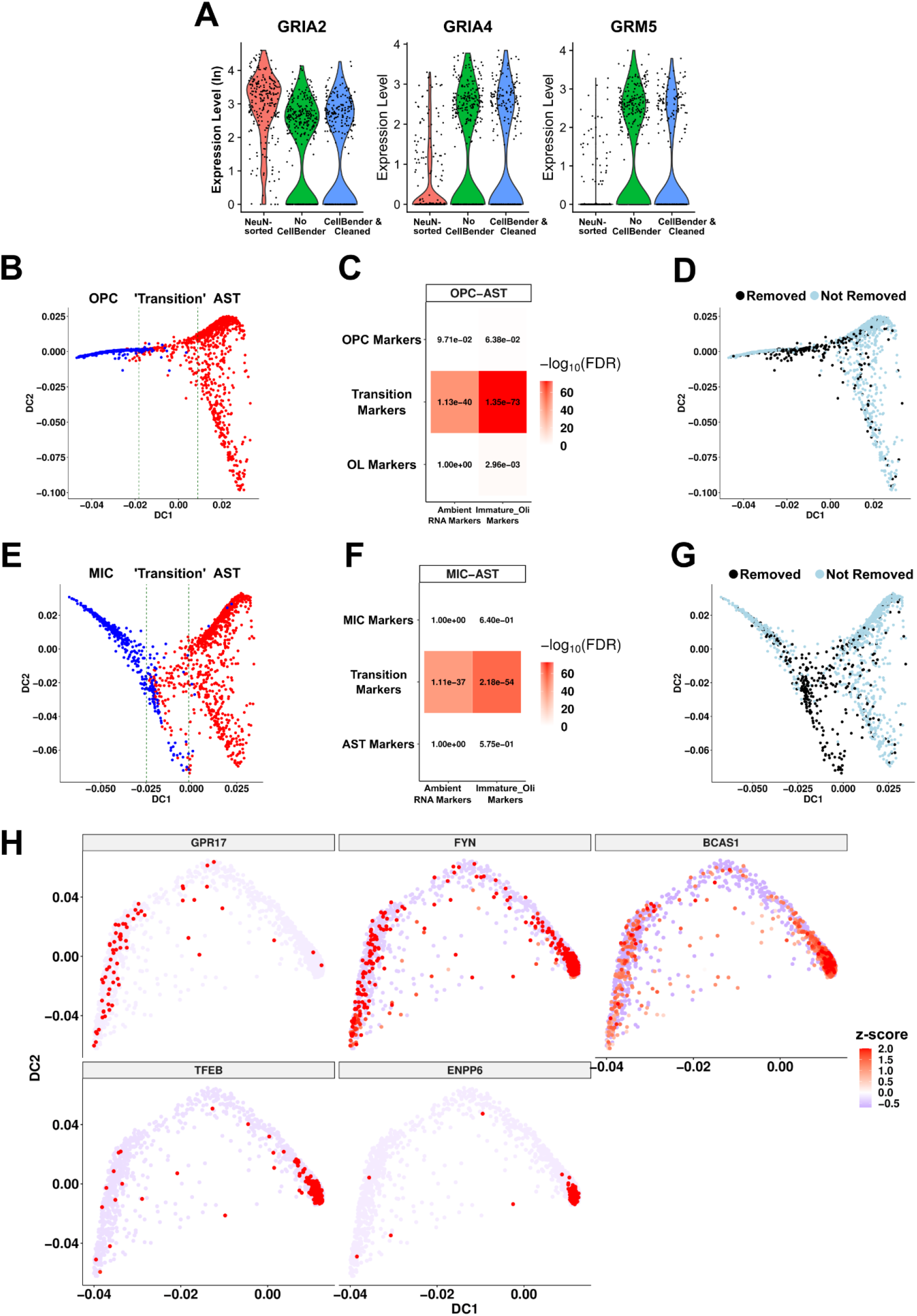
Immature oligodendrocytes are explained by ambient RNA contamination. **(A)** Violin plots of glutamatergic receptors in NeuN-sorted OPCs and SD1OPCs with or without ambient RNA removal. **(B)** Pseudotime trajectory as reconstructed with *destiny* between OPC and AST (astrocyte). ‘Transition’ zone was determined as 400 cells around the middle cell based on DC1. **(C)** Enrichments between trajectory zones (OPC, Transition, AST) and ambient RNA or immature oligodendrocyte markers (Fisher’s exact test, numbers indicate FDR, color scale is −log_10_(FDR). **(D)** Same lineage trajectory as **(B)** with cells removed after subcluster cleaning highlighted. **(E-G)** Same as OPC-AST from **(B-D)** but between MIC (microglia) and AST. **(H)** Z-transformed gene expression of COP or Pre-OL (premyelinating oligodendrocyte) markers in OPC-OL lineage trajectory.

**Extended Figure 6.**
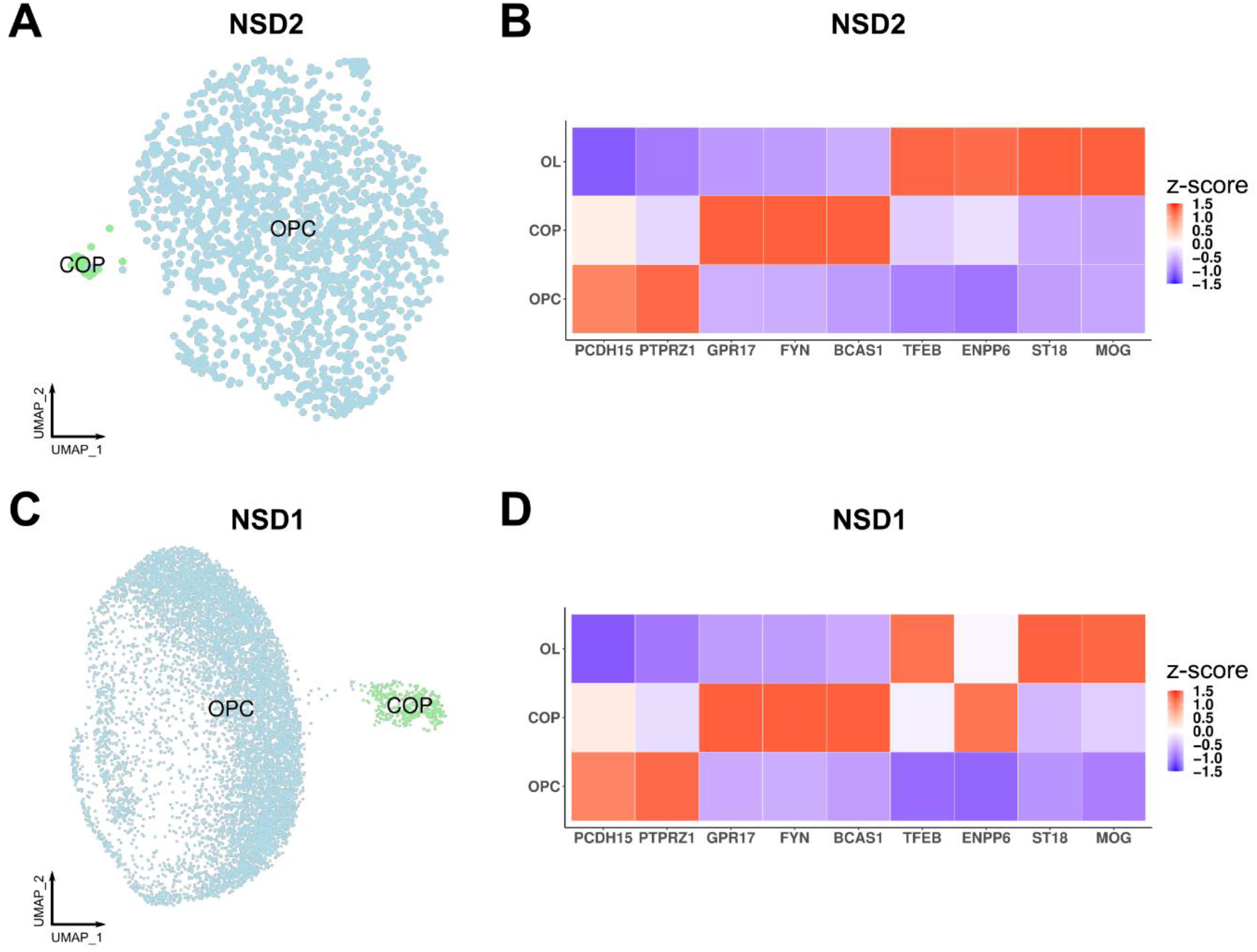
Committed progenitor cells in other adult human brain snRNA-seq datasets. **(A, C)** UMAP of OPC subclustering in **(A)** NSD2 and **(C)** NSD1 datasets. COP: committed OPCs. **(B, D)** Heatmap of oligodendrocyte lineage markers (z-scored across cell types per marker gene) in **(B)** NSD2 and **(D)** NSD1 datasets.

**Extended Figure 7.**
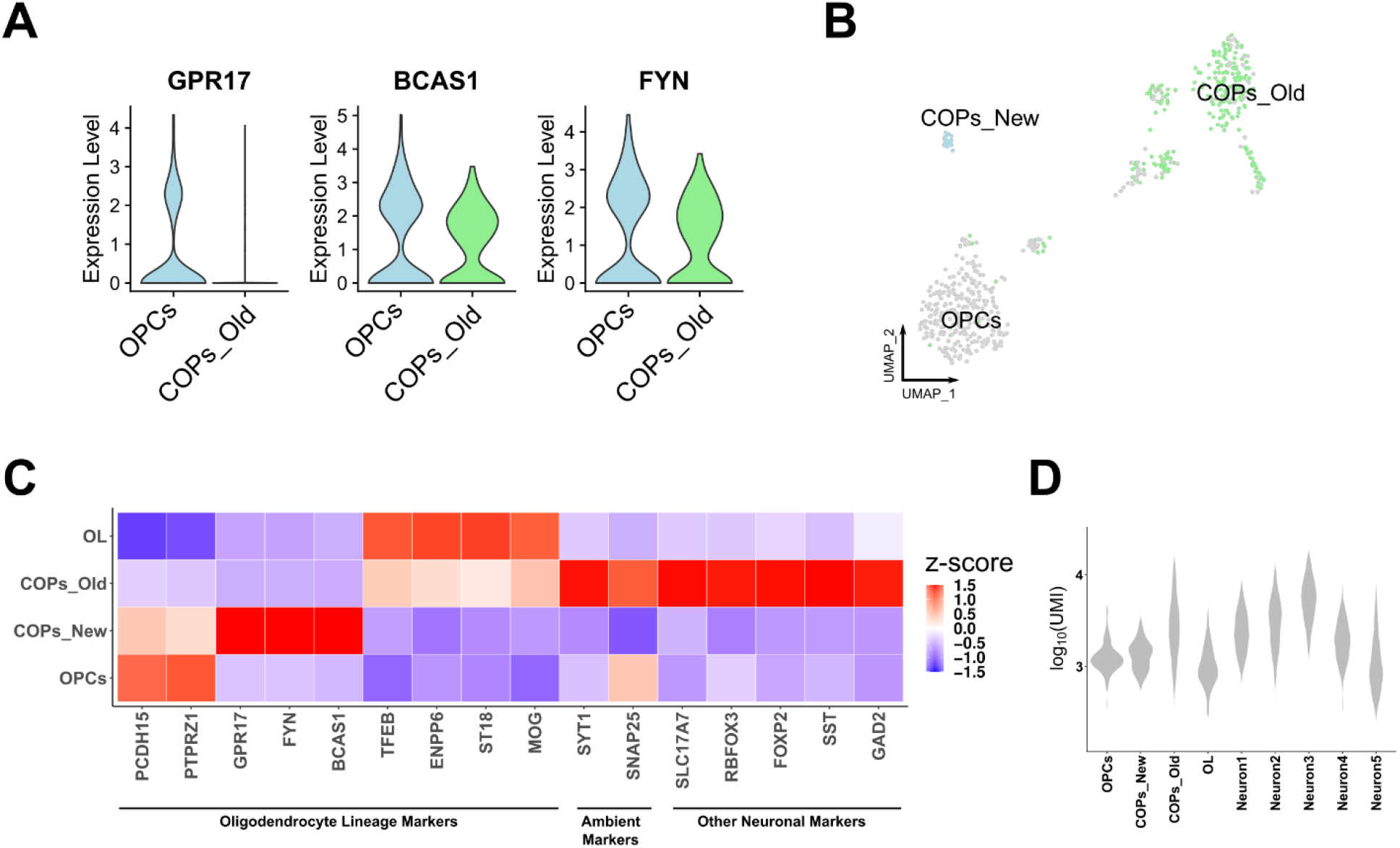
Re-assessment of previous COP annotation in human brain snRNA-seq dataset from white matter. **(A)** Expression levels (normalized, log transformed) of COP markers in the original annotation of OPCs and COPs. **(B)** UMAP plot of OPCs and COPs. The small subpopulation suspected to be real COPs is shown as ‘COPs_New’ whereas the cells previously annotated as COPs are shown as ‘COPs_Old’. **(C)** Heatmap of oligodendrocyte lineage markers, two top ambient RNA markers (*SYT1*: Nuclear, *SNAP25*: Extra-nuclear) and other neuronal markers in the same dataset. Colors indicate z-scored expression across cell types per marker gene. **(D)** Log_10_ transformed UMI count values per cell type. OL: oligodendrocytes. Neuronal cell types are annotated as in the original publication^13^.

**Extended Figure 8.**
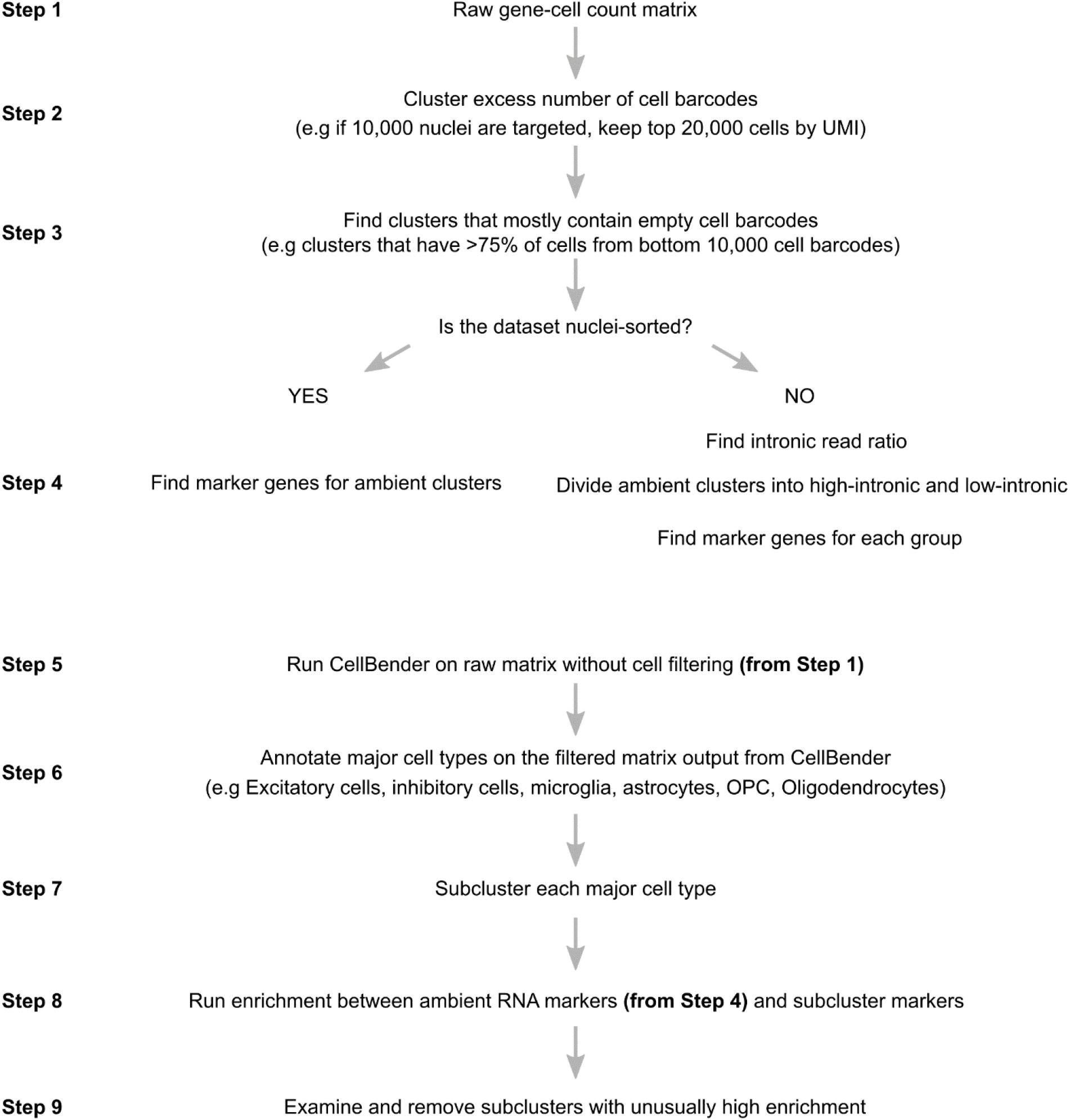
Stepwise guideline of ambient RNA marker detection and ambient RNA removal. Steps 1-4 describes how to identify ambient RNA markers in the given dataset. Steps 5-9 describes how to use this information to further remove ambient RNA contaminated cell barcodes after a formal ambient RNA contamination removal tool such as CellBender.

